# Cannabinoid agonist WIN55,212-2 prevents scopolamine-induced impairment of spatial memory

**DOI:** 10.1101/2024.12.12.628159

**Authors:** Marta Moreno-Rodríguez, Iker Bengoetxea de Tena, Jonatan Martínez-Gardeazabal, Gorka Pereira-Castelo, Alberto Llorente-Ovejero, Iván Manuel, Rafael Rodríguez-Puertas

## Abstract

The endocannabinoid system is involved in diverse processes, like learning and memory, governed by cholinergic neurotransmission. Recent research demonstrates that in a model of dementia derived from basal forebrain cholinergic degeneration, WIN55,212-2 improves cognition through increased cortical choline levels. However, the effect of cannabinoids on cholinergic deficits is still under investigation. In this work, we studied the effect of this treatment in a pharmacological model of transient cholinergic hypofunction by the acute administration of the muscarinic antagonist, scopolamine, in spatial, recognition and aversive memory tests. Scopolamine induced memory impairment was observed in the three tests and, importantly, the cannabinoid subchronic treatment with low doses of WIN55,212-2 prevented this deleterious effect in spatial memory when evaluated in spatial Barnes maze test. Autoradiographic studies indicate that, following the WIN55,212-2 treatment, cannabinoid receptor density increased in the motor and somatosensory cortices. In layers I-V of the motor cortex, the activity of cannabinoid and muscarinic receptors also increased. These results suggest that WIN55,212-2, through the activation of CB_1_ receptors, indirectly elevates the muscarinic tone in key cortical areas for learning and memory, preventing the memory deficits induced by scopolamine specifically in spatial memory. This highlights the importance of the crosstalk between the endocannabinoid and the cholinergic system for learning and memory processes and suggest that cannabinoid agonists might be an alternative for the treatment of cognitive deficits associated with cholinergic dysfunction.

## 1. Introduction

The endocannabinoid system (eCB), comprising cannabinoid receptors, CB_1_ and CB_2_, endogenous ligands and metabolic enzymes, regulates relevant physiological processes including pain, appetite, immune responses and cognitive processes (Lu & Mackie, 2021). These functions are predominantly regulated by CB_1_ receptors, which are among the most expressed and active G protein-coupled receptors (GPCRs) in the central nervous system (CNS) (Martinez Ramirez et al., 2023).

Numerous studies indicate that cannabinoids negatively affect learning and memory in physiologically normal states (Urits et al., 2021) e.g., Δ-9-tetrahydrocannabinol (Δ^9^-THC) consumption has been linked to impairments in various brain functions, including catalepsy and acute disruptions in attention and cognitive task performance (Stella, 2023). However, growing evidence suggests that the effects of cannabinoids are biphasic and dose-dependent, and are influenced by the preexisting cognitive status. In fact, lower doses of cannabinoids might be beneficial in some contexts, including aging, Alzheimer’s disease (AD) and other dementia-related neurodegenerative conditions (Bilkei-Gorzo et al., 2017; Ozaita & Aso, 2017; Llorente-Ovejero et al., 2018; for review see Pereira-Castelo et al., 2024).

Some of these effects result from interactions between the eCB system and other neurotransmitters in the CNS, such as the cholinergic system, which is essential for memory formation (Gedankien et al., 2023). As an example of these interactions, presynaptic CB_1_ receptors at the medial habenula regulate the expression of aversive memories through the specific control of cholinergic neurotransmission (Soria-Gómez et al., 2015). In a spatial memory task, the cognitive impairment produced by a treatment with the cannabinoid receptors agonist WIN55,212-2 was mediated by cholinergic hypofunction (Robinson et al., 2010) and, notably, both CB_1_ and M_2_ receptors are preferentially presynaptic and coupled to inhibitory G proteins (Nyíri et al., 2005). In this line, a previous study from our group investigated the effect of a subchronic WIN55,212-2 treatment on learning and memory in a model of basal forebrain cholinergic hypofunction, and the observed cognitive improvement was mediated by increased levels of acetylcholine (ACh) together with the restoration of choline-containing cortical lipids (Moreno-Rodríguez et al., 2024).

Various studies performed using the muscarinic antagonist scopolamine further support the interaction between the cholinergic and the eCB systems, although data concerning the impact of cannabinoid compounds on scopolamine-induced effects is inconclusive. In fact, studies performed with inverse agonists of CB_1_ receptors report both a potentiation of the disruptive effects of scopolamine (Nakamura-Palacios et al., 2000), or improved cognitive performance following administration (Dillon et al., 2011). In a passive avoidance task, bilateral microinjection at the basolateral amygdala (BLA) of arachydonilcyclopropylamide (ACPA), a CB_1_ receptor agonist, improved scopolamine-induced memory impairment, while co-administration of ineffective scopolamine doses with AM251, a cannabinoid CB_1_ receptor antagonist, mimicked the amnesic effect obtained with higher scopolamine doses (Nedaei et al., 2016).

On this basis, the present study evaluated, *in vivo*, the effect of a cannabinoid treatment with low doses of WIN55,212-2 in preventing the transient amnesia induced by acute scopolamine administration in paradigms related to spatial, recognition and aversive memory, and analyzed the neurochemical correlates behind the observed behaviors using a pharmacological approach by autoradiographic assays.

## 2. Material and methods

### 2.1 Reagents and drugs

All the compounds necessary for the different procedures were of the highest quality commercially available for the purpose of our studies.

[^3^H]CP55,940 (149 Ci/mmol) and [^35^S]GTPγS (1250 Ci/mmol) were acquired from Revvity (Waltham, MA, USA). The [^3^H] microscales and [^14^C] microscales used as standards in the autoradiographic experiments were purchased from ARC (American Radiolabeled Chemicals, Saint Louis, MO, USA). The β-radiation sensitive films, Kodak Biomax MR, bovine serum albumin (BSA), DL-dithiothreitol (DTT), guanosine 5’-diphosphate (GDP), guanosine 5’-O-3-thiotriphosphate (GTPγS), ketamine and xylazine were acquired from Sigma-Aldrich (St Louis, MO, USA).

2-carbamoyloxyethyl-trimethyl-azanium (Carbachol) and (α,S)-α- (Hydroxymethyl)benzeneacetic acid (1α,2β,4β,5α,7β)-9-methyl-3-oxa-9-azatricyclo[3.3.1.02,4]non-7-yl ester hydrobromide (Scopolamine) were acquired from Sigma-Aldrich (St Louis, MO, USA). (11R)-2-Methyl-11-[(morpholin-4-yl)methyl]-3-(naphthalene-1-carbonyl)-9-oxa-1 azatricyclo[6.3.1.04,12]dodeca-2,4(12),5,7-tetraene (WIN55,212-2) was acquired from Tocris (Bristol, UK).

### 2.2 Animals

Every effort was made to minimize animal suffering and to use the minimum number of animals possible throughout the whole study. All procedures were performed in accordance with European animal research laws (Directive 2010/63/EU) and the Spanish National protocols, and were approved by the Local Ethical Committee for Animal Research of the University of the Basque Country (CEEA M20-2018-52 and 54 and M20/2023/352).

Male Sprague-Dawley rats (n = 90) weighing 200-300 g were housed in groups of 3-4 per cage at a temperature of 22°C and in a humidity-controlled (65%) room with a 12:12 hours light/dark cycle, with access to food and water *ad libitum*. These rats were administered with scopolamine/vehicle and WIN55,212-2/vehicle and were used for the performance of the different behavioral tests. Some of the brains were also used for autoradiographic studies (n = 35).

### 2.3 Behavioral tests

#### 2.3.1 Barnes maze test

The test was conducted on a white circular platform (130 cm diameter, 1 m above the floor) with 20 equally spaced holes. Only one hole led to a dark escape chamber beneath the platform. Bright lights placed around the platform created an unpleasant atmosphere, encouraging rats to find the target hole. Visual cues on the walls helped with spatial orientation.

The test had two phases over five days: acquisition (4 days) and probe trial (day 5). During acquisition, rats were placed on the platform, given 3 minutes to find the target hole, and were guided to it if they failed. Four trials were conducted daily with 15 min of inter-trial intervals. On the fifth day, the day after the last acquisition trial, a single 90 s probe trial was performed, where memory retention was tested without the escape box present. In this phase, the platform was virtually divided into four quadrants and the time spent by the rats in the target quadrant, where the escape box was previously located, was measured as indicative of spatial memory. An automated tracking system (SMART, Panlab S.L., Barcelona, Spain) recorded latency, path length, and speed during the acquisition phase, while quadrant time was used to assess spatial memory during the probe trial.

#### 2.3.2 Novel object recognition test

The test was conducted in a white open-field arena (90 × 90 × 50 cm) (Panlab S.L., Barcelona, Spain) under low light. It included four phases over five days: habituation (3 days), familiarization, short-term testing (5 hours later), and long-term testing (24 hours later).

Before each phase, rats were handled gently for 1 minute. During habituation phase, rats explored the arena for 5 minutes. In the familiarization phase, two identical objects (object A) were placed in opposite corners of the arena. Rats explored them until a combined 25-second exploration threshold was reached. In short-term testing, rats were presented with the familiar object (A) and a new object (B) for 5 min. Long-term testing, conducted 24 hours after the familiarization phase, involved the familiar object (A) and another new object (C). A video camera recorded rat behavior, and exploration times for both familiar and novel objects were measured as indicative of recognition memory. The object discrimination ratio (DR) was calculated, with scores near zero indicating no preference, negative scores indicating a preference for the familiar object, and positive scores indicating preference for the novel object.

#### 2.3.3 Passive avoidance test

The behavioral test was conducted in a shuttle box (PanLab S.L. Barcelona, Spain) with two compartments: a larger, brightly lit, white compartment (31 × 31 × 24 cm) and a smaller, dark, black compartment (19.5 × 10.8 × 12 cm), separated by a guillotine door. The test had two phases: acquisition and retention. Before each phase, rats were habituated to the experimental room. In the acquisition phase (day 1), rats explored the white compartment for 30 seconds before the door to the black compartment opened. Upon instinctively crossing into the black compartment, the door closed, and a mild foot shock (0.4 mA for 2 seconds) was administered. Rats remained in the black compartment for 15 seconds before being returned to their cage. The shuttle box was sanitized with ethanol (70%) between trials. Rats that did not cross into the black compartment in this phase were excluded from the test. In the retention phase (day 2), rats were again placed in the white compartment, and after 30 s, the door to the black compartment opened. Rats had 5 min to decide whether to cross; those that crossed received no shock. The behavior of the rats was recorded with a video camera positioned above the shuttle box, with acquisition latency (day 1) and step-through latency (day 2) measured. Longer latencies in the retention phase indicated a passive avoidance response, reflecting positive performance.

#### 2.3.4 Hot plate test

The test was performed in a hot plate apparatus (Leica Biosystems, Barcelona, Spain) consisting in a cylindrical see-through Plexiglas wall (19 x 30 cm) located above a plate. The plate warms through an electric resistance and is equipped with a timer and a thermostat. On the top of the cylinder, a metal grid is located to hold the rat when it jumps.

Rats were placed on the metal plate, which was previously warmed (55 ± 0.5 °C). The time spent by the rats until jumping was recorded (jump latency). The latency to paw-licking was also measured. Both parameters, following a noxious thermal stimulus, were used as indicative of nociceptive threshold.

#### 2.3.5 Electrical shock evoked pain threshold

The procedure was performed in the same shuttle box used for the PA test. This apparatus has a grid floor and electrical foot shocks of different potency can be delivered.

Rats were placed in the white compartment and received mild foot shocks, beginning at 0.0 mA, gradually increasing by 0.05 mA, until the first vocalization was measured, which is an indicative of nociception and discomfort.

### 2.4 WIN55,212-2 administration in a pharmacological model of muscarinic antagonism

Scopolamine was dissolved in saline 0.9 % and was administered intraperitoneally (2 mg/kg) in a volume of 10 ml/kg, 30 min before the performance of the PA test acquisition phase, the NORT test short and long-term testing phases and the BM probe trial phase. Control group received vehicle without scopolamine.

For the evaluation of the effect of a cannabinoid treatment in the amnesic effects elicited by scopolamine, WIN55,212-2 was intraperitoneally administered once daily (0.5 mg/kg), one hour before every phase of each test. WIN55,212-2 was dissolved in pure DMSO and diluted with kolliphor and 0.9 % saline, in a 1:1:18 proportion. Control group received vehicle without WIN55,212-2. Rats were randomly divided into four groups for each test: vehicle (VEH), WIN55,212-2 (WIN), scopolamine (SCOP) and WIN55,212-2 + scopolamine (WIN+SCOP).

### 2.5 Autoradiographic studies

#### 2.5.1 Functional [^35^S]GTPγS autoradiography

For the performance of functional autoradiography to study muscarinic and cannabinoid receptors, brain sections were air dried for 30 min, followed by two consecutive incubations of 30 min each in an HEPES-based buffer (50 mM HEPES, 100 mM NaCl, 3 mM MgCl_2_, 0.2 mM EGTA and 0.5% BSA, pH 7.4) at 30°C to remove the endogenous ligands. Slices were then incubated for 2 h at 30°C in the same buffer supplemented with 2 mM GDP, 1 mM DTT and 0.04 nM [^35^S]GTPγS. Basal binding was determined in two consecutive slices in the absence of the agonist. The agonist-stimulated binding was determined in another consecutive slice in the presence of the corresponding receptor agonists, carbachol (10 µM) for muscarinic receptors and WIN55,212-2 (10 µM) for cannabinoid receptors. Non-specific binding was defined by competition with unlabeled GTPγS (10 µM) in another section. Tissue slices were finally washed twice in cold (4°C) HEPES (50 mM) buffer (pH 7.4), dried and exposed for 48 h to β-radiation sensitive films with a set of [^14^C] standards calibrated for [^35^S]. Calibrated films were scanned and quantified using Fiji software. Data was expressed as the % of stimulation over basal.

#### 2.5.2 [^3^H]CP55,940 receptor autoradiography

For the performance of cannabinoid receptors autoradiography, brain sections were air dried for 30 min and then immersed in Coplin jars for preincubation in a buffer containing 50 mM Tris-HCl and 1% of BSA (pH 7.4) for 30 min, at room temperature, to remove endogenous ligands. Tissue slices were later incubated in the presence of the [^3^H]CP55,940 radioligand (3 nM) for 2 h at 37°C. Non-specific binding was measured by competition with non-labeled CP55,940 (10 µM) in another consecutive slice. Following the incubation, tissue slices were washed with an ice-cold preincubation buffer, dipped in distilled water and dried overnight. To generate autoradiograms, dry sections were exposed to β-radiation-sensitive films in hermetically closed cassettes for 21 days at 4°C. For the calibration of the optical densities to fmol/mg tissue equivalent, [^3^H] microscales were exposed to the films. Calibrated films were scanned and quantified using Fiji software (Fiji, Bethesda, MD, USA). Data was expressed as fmol/g of tissue equivalent (t.e.).

### 2.6 Statistical analysis

Data from behavioral tests was analyzed using a Mann-Whitney test for two groups and Kruskal-Wallis test followed by Dunn’s *post hoc* test for more than two groups. Repeated measures two-way ANOVA followed by *post-hoc* test Bonferroni’s was used for repeated measures of the same parameter in different days (total latency, total path length and speed in BM test). Step-through latency times of PA test were represented as Kaplan-Meier survival curves and analyzed using a log-rank/Mantel–Cox test, as previously described by our group (Llorente-Ovejero et al., 2017; Bengoetxea de Tena et al., 2022). Data from autoradiographic assays was analyzed using a Mann-Whitney test. The threshold for statistical significance was set at p < 0.05. Statistical analyses and data representation were performed using GraphPad Prism 9 (GraphPad Software, Boston, MA, USA).

## 3. Results

### 3.1 WIN55,212-2 prevents scopolamine-induced amnesia in a spatial memory test

BM test was performed to evaluate the effect of WIN55,212-2 in preventing scopolamine-induced impairment of spatial memory. Throughout the four days of acquisition, both WIN55,212-2 and vehicle-treated groups decreased the total latency and the total path length (Fig. 1A and Fig. 1B). On the first trial, this parameter was significantly higher in the WIN55,212-2-treated group (VEH *vs*. WIN, p < 0.05; Fig. 1A), indicating a slight delay in the learning curve. The average speed increased in each trial for both groups, but WIN55,212-2-treated rats walked more slowly than vehicle-treated ones overall (VEH *vs*. WIN, p < 0.05; Fig. 1C).

**Fig. 1.**
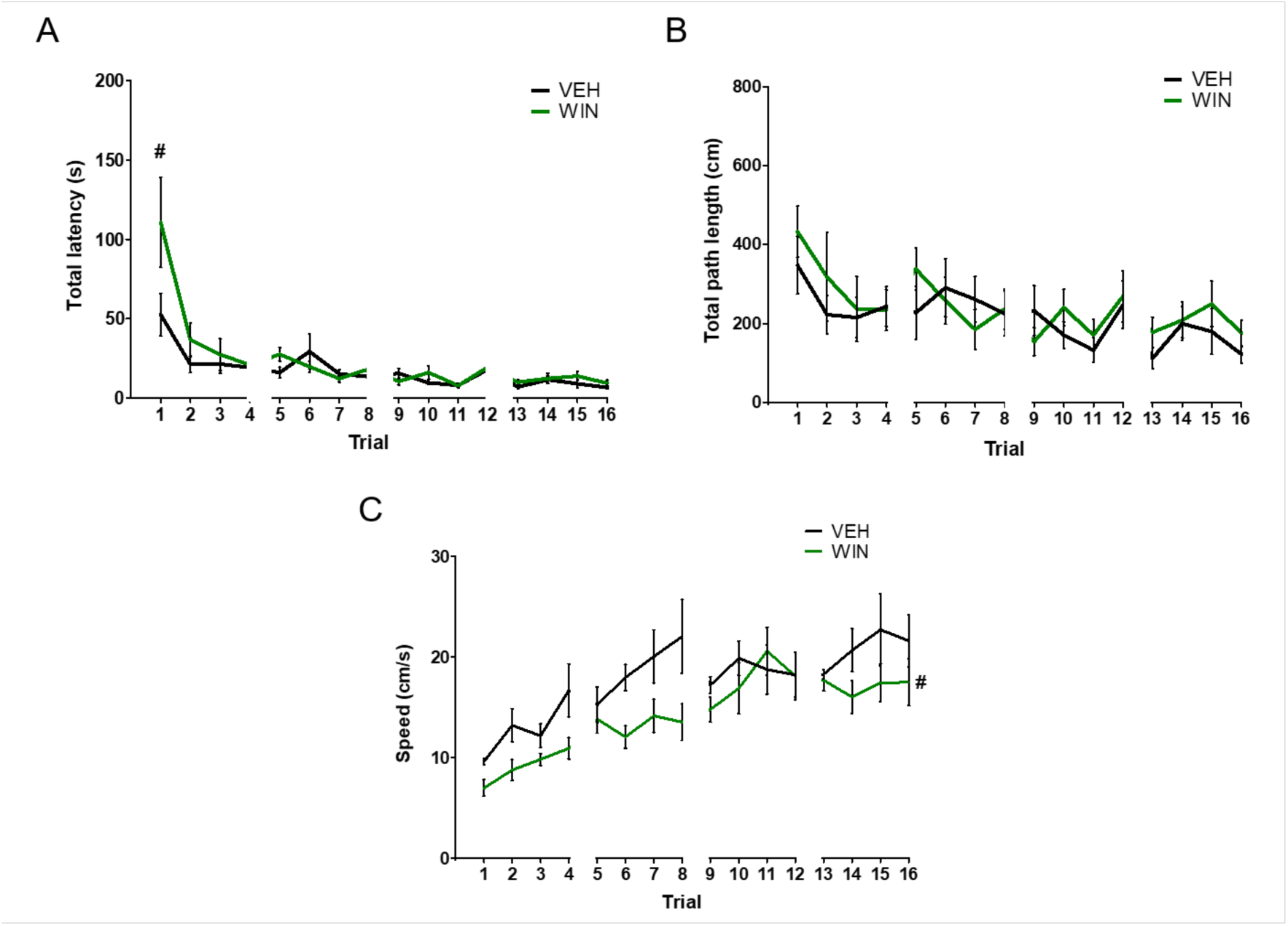
Effect of WIN55,212-2 on locomotion and learning in BM test. (A) Total latency during the four days of learning of BM test. No significant differences were observed between VEH and WIN groups comparing both curves, but significant differences were observed between the two groups in the first trial (Repeated measures two-way ANOVA, *post-hoc* test Bonferroni’s, ^#^p < 0.05, VEH *vs*. WIN). (B) Total path length during the four days of learning of BM test. No significant differences were observed between VEH and WIN groups comparing both curves (Repeated measures two-way ANOVA, *post-hoc* test Bonferroni’s). (C) Mean speed during the four days of learning of BM test. Significant differences were observed between VEH and WIN groups comparing both curves (Repeated measures two-way ANOVA, *post-hoc* test Bonferroni’s, ^#^p < 0.05, VEH *vs*. WIN).

On probe trial, after the last training day, the latency in the target quadrant was measured as an indicator of spatial memory in all groups. The administration of WIN55,212-2 to control rats did not produce an impairment in spatial memory (VEH *vs*. WIN, p > 0.05; Fig. 2A). Scopolamine-treated rats spent a significantly lower time in the target quadrant (VEH *vs*. SCOP, p < 0.001; Fig. 2A), indicating spatial memory deficits following its administration. Importantly, a subchronic treatment with WIN55,212-2 prevented the amnesic effect exerted by scopolamine in BM test (SCOP *vs*. SCOP+WIN, p < 0.01; Fig. 2A).

**Fig. 2.**
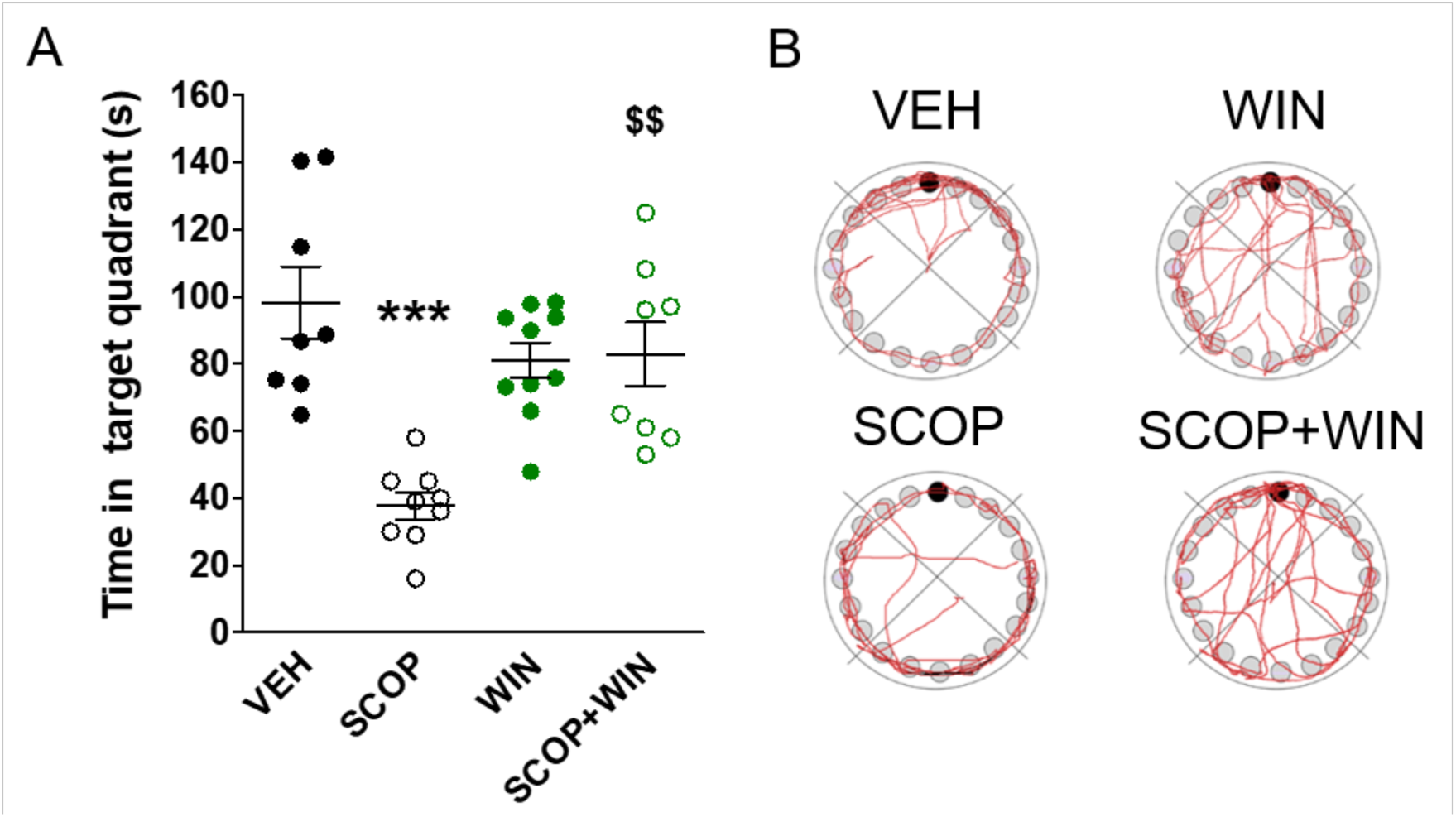
WIN55,212-2 prevents the scopolamine-induced impairment of spatial memory. (A) Time spent in the target quadrant on the probe trial day of BM test (Kruskal–Wallis test, *post-hoc* test Dunn’s multiple comparison, ***p < 0.001 VEH *vs*. SCOP, ^$$^p < 0.01 SCOP *vs*. SCOP+WIN). (B) Representative trajectories of rats from each group during 180 s in the probe trial day of BM test. Note the accumulation of trajectories in the target quadrant (where the target hole is located, depicted in black in the figure) for VEH, WIN and SCOP+WIN groups, as opposed to SCOP group.

Considering the positive effects of a cannabinoid treatment in BM, we analyzed its effect on preventing the amnesic effects induced by scopolamine in other memory tests, including the novel object recognition test (NORT) and passive avoidance (PA) test.

### 3.2 Effect of a cannabinoid treatment on a pharmacological model of muscarinic antagonism in recognition memory

Recognition memory was evaluated at two different time points, short (5 h post-familiarization) and long-term (24 h post-familiarization). In the short-term test, the treatment with WIN55,212-2 impaired memory in control rats (VEH *vs*. WIN, p < 0.001; Fig. 3A). Notably, the administration of scopolamine did not significantly decrease the DR (VEH *vs*. SCOP, p > 0.05; Fig. 3A).

**Fig. 3.**
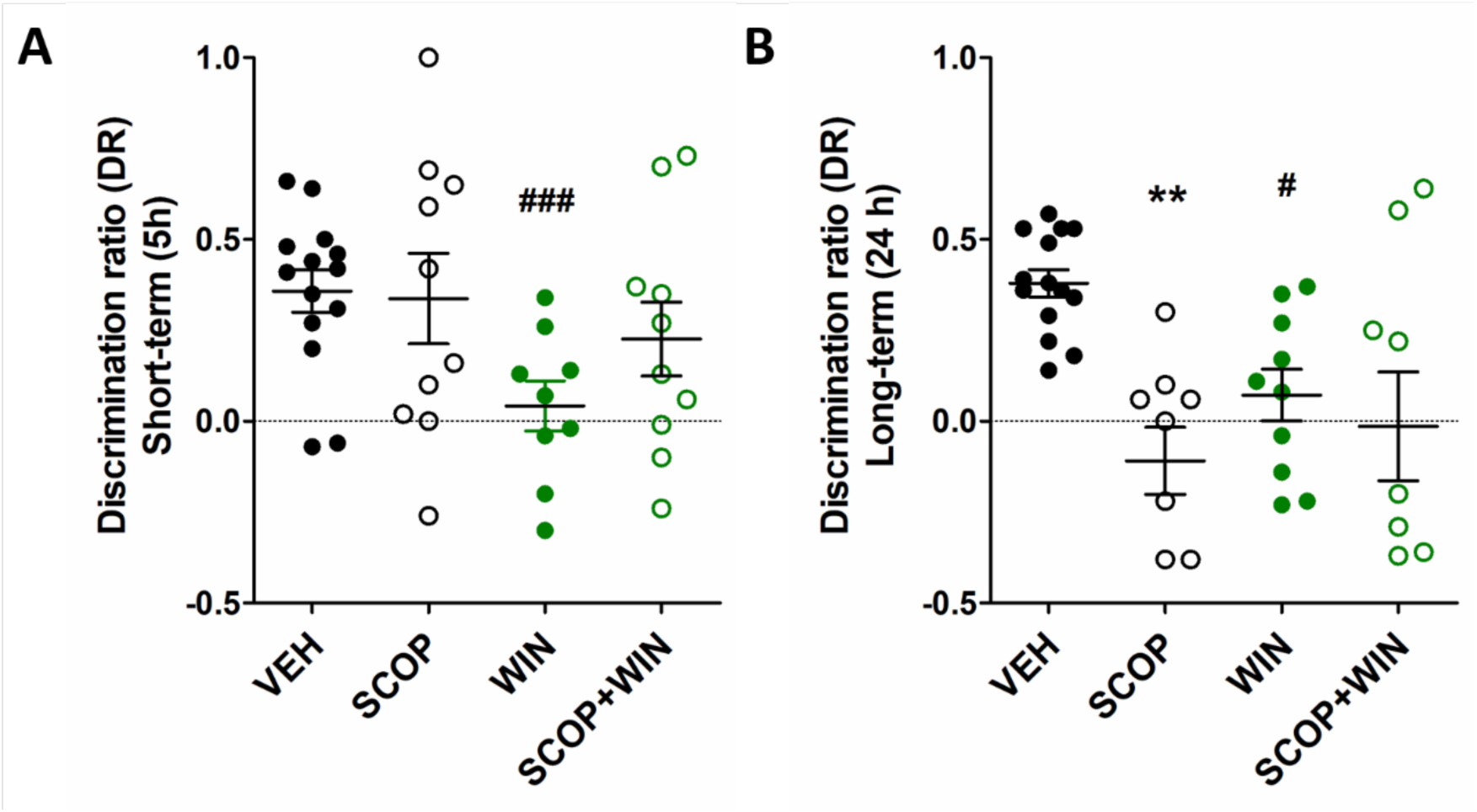
Effect of WIN55,212-2 in a model of muscarinic antagonism in recognition memory in the short-term and the long-term. (A) Discrimination ratio in the short-term testing phase of NORT (Kruskal–Wallis test, *post-hoc* test Dunn’s multiple comparison, ^###^p < 0.001 VEH *vs*. WIN). (B) Discrimination ratio in the long-term testing phase of NORT (Kruskal–Wallis test, *post-hoc* test Dunn’s multiple comparison, **p < 0.01 VEH *vs*. SCOP, ^#^p < 0.05 VEH *vs*. WIN).

In the long-term, the subchronic treatment with WIN55,212-2 also showed a deleterious effect on memory in control rats (VEH *vs*. WIN, p < 0.05; Fig. 3B). In contrast to what was observed in the short-term test, the administration of scopolamine also caused a significant decrease in the DR (VEH *vs*. SCOP, p < 0.01; Fig. 3B), indicating memory impairment following the administration of a muscarinic antagonist in long-term recognition and working memory, but not in short-term one. The administration of WIN55,212-2 to rats that also received scopolamine only produced a slight increase in the DR and did not prevent or reverse the observed memory deficits (SCOP *vs*. SCOP+WIN, p > 0.05; Fig. 3B).

### 3.3 Effect of a cannabinoid treatment on a pharmacological model of muscarinic antagonism in an aversive memory test

The administration of scopolamine impaired both acquisition latency (VEH *vs*. SCOP, p < 0.01, Fig. 4A) and step-through latency time (VEH *vs*. SCOP, p < 0.001, Fig. 4B), provoking transient memory deficits. The cannabinoid treatment with WIN55,212-2 alone did not affect acquisition latency, but decreased the step-through latency time (VEH *vs*. WIN, p < 0.01, Fig. 4B). The subchronic administration of WIN55,212-2 did not prevent the amnesic effect of scopolamine observed in the step-through latency time (SCOP *vs*. SCOP+WIN, p > 0.05, Fig. 4B).

**Fig. 4.**
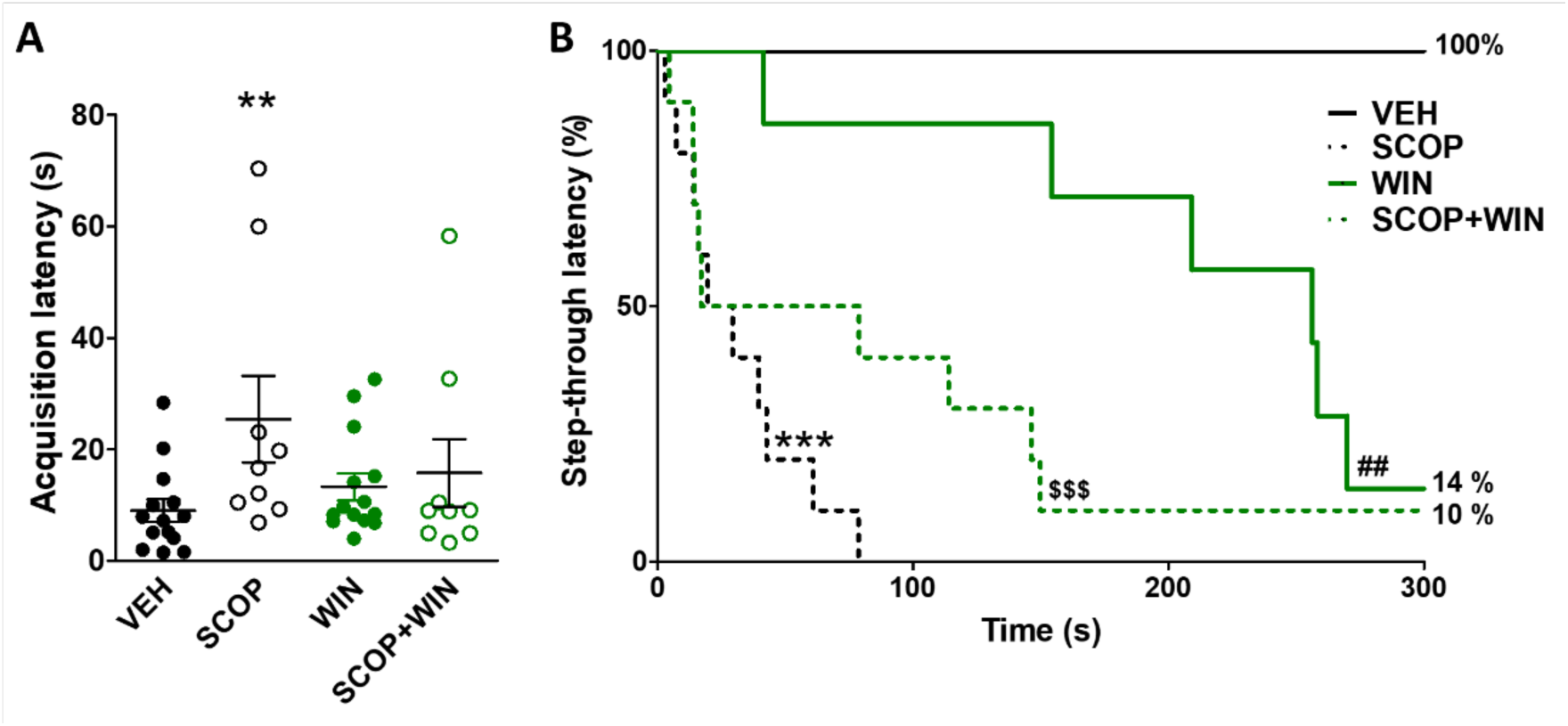
Treatment with WIN55,212-2 in a model of muscarinic antagonism evaluated in PA test. (A) Acquisition latency times in the learning trial of the PA test (Kruskal–Wallis test, *post-hoc* test Dunn’s multiple comparison, **p < 0.01, VEH *vs*. SCOP). (B) Step-through latency times of PA test represented as Kaplan-Meier survival curves (log-rank/Mantel–Cox test, ***p < 0.001 VEH *vs*. SCOP, ^##^p < 0.01 VEH *vs*. WIN, ^$$$^p < 0.001 VEH *vs*. SCOP+WIN). The median latency was 24.55 s for SCOP group, 256.00 s for WIN group and 48.00 s for SCOP+WIN group.

Considering the analgesic effects of cannabinoid agonists (Manzanares et al., 2006), we hypothesized that decreased pain sensitivity following the WIN55,212-2 administration could be a bias in the effect of the treatment observed in PA test. Thus, we measured the effect of this treatment on pain response using the hot plate test. We observed that WIN55,212-2 treatment produced analgesia, as indicated by the increases the jump latency (VEH *vs*. WIN, p < 0.05, Fig. 5A) and the latency to licked paw (VEH *vs*. WIN, p < 0.01, Fig. 5B). We also measured pain response using a protocol designed to evaluate nociception produced by an electrical shock in PA test, the electrical shock evoked pain threshold, and we observed that the group of rats treated with WIN55,212-2 required a higher shock intensity to evoke a first vocalization (VEH 0.0375 ± 0.0082 mA *vs*. WIN 0.0714 ± 0.0101 mA, p < 0.01), also indicating an analgesic effect of the cannabinoid treatment. These results indicate a possible bias of the results obtained in PA test following the WIN55,212-2 administration.

**Fig. 5.**
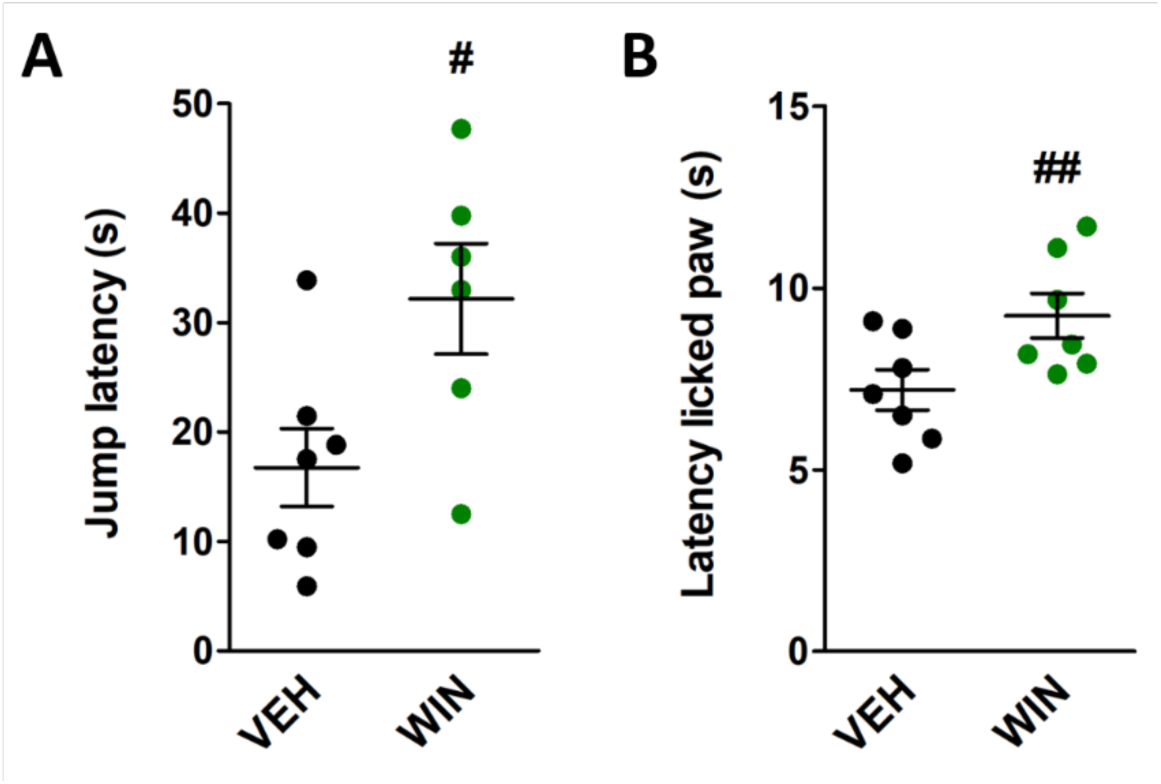
Analgesic effect of WIN55,212-2 in the hot plate test. (A) Jump latency in the hot plate test (Mann Whitney test, ^#^p < 0.05 VEH *vs*. WIN). (B) Latency to licked paw in the hot plate test (Mann Whitney test, ^##^p < 0.01 VEH *vs*. WIN).

### 3.4 The subchronic WIN55,212-2 administration altered muscarinic and cannabinoid receptor activity

The effect of WIN55,212-2 administration on muscarinic and cannabinoid receptors was studied to elucidate the neurochemical correlates behind the observed behaviors. The density and activity of muscarinic and cannabinoid receptors was analyzed by means of autoradiography in rats that had performed BM test, given the positive effect exerted by the treatment in this test. Considering that scopolamine was administered acutely and given its fast excretion and low bioavailability (Tian et al., 2015), the autoradiography data only represents the effect of the WIN55,212-2 treatment.

The [^35^S]GTPγS binding stimulated by carbachol, an agonist of muscarinic receptors, was measured in brain areas related to learning and memory to localize and determine the activity of G_i/o_ proteins coupled to M_2_/M_4_ receptors (Fig. 6). No differences were observed between the groups treated with vehicle and WIN55,212-2 in basal binding, i.e, [^35^S]GTPγS binding in the absence of the agonist. The G_i/o_-coupled M_2_/M_4_ receptor activity induced by carbachol was increased in the WIN55,212-2-treated group in layers I-V of the motor cortex (VEH *vs*. WIN, p < 0.05; Fig. 6, Table 1), in some of the septal nuclei, the medial septum and the horizontal diagonal band (VEH *vs*. WIN, p < 0.05; Fig. 6, Table 1), and in the pyramidal layer of the CA3 region of the hippocampus (VEH *vs*. WIN, p < 0.05; Fig. 6, Table 1).

**Fig. 6.**
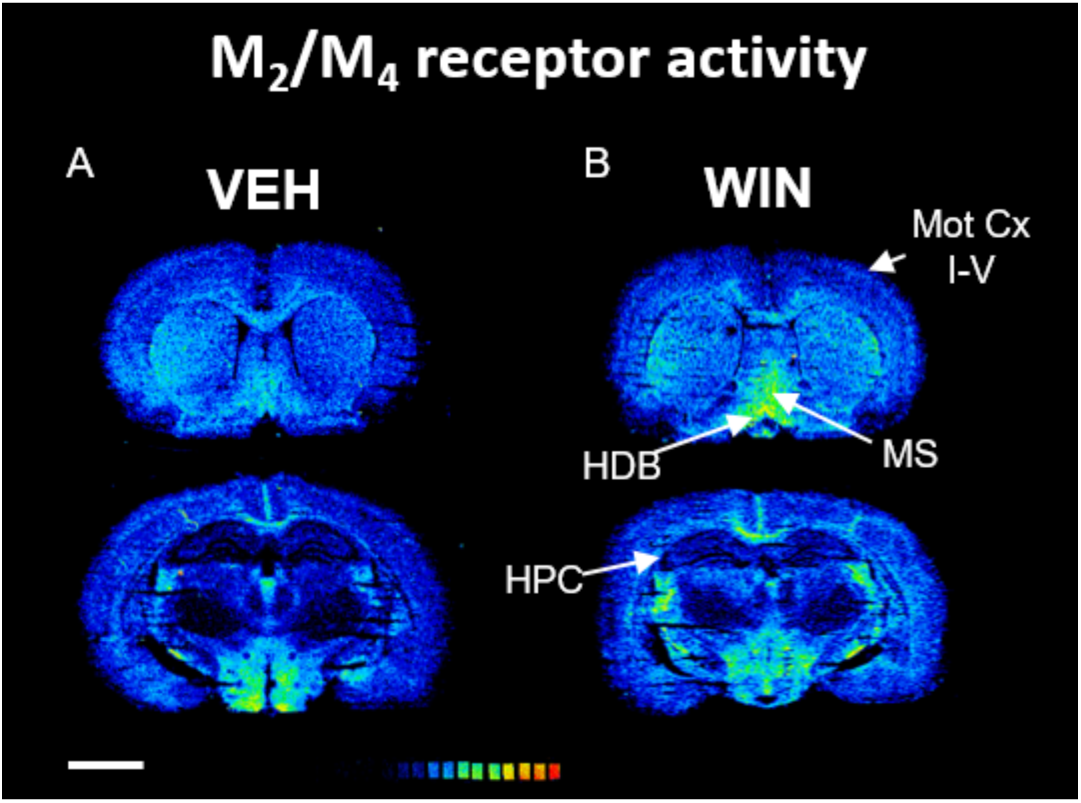
Representative autoradiograms showing the activity of M2/M4 receptors. Coronal sections corresponding to (A) VEH and (B) WIN groups showing [^35^S]GTPγS binding stimulated by carbachol in the horizontal diagonal band (HDB), layers I-V of the motor cortex (Mot Cx I-V), medial septum (MS) and the hippocampus (HPC). Scale bar: 4 mm.

**Table 1.** [^35^S]GTPγS binding of carbachol-stimulated and WIN55,212-2-stimulated receptor activity expressed as the percentage of stimulation over basal, in different brain areas related to learning and memory.

| Brain region | Carbachol stimulation<br>(% over basal) |  | WIN55,212-2 stimulation<br>(% over basal) |  |
| --- | --- | --- | --- | --- |
|  | VEH | WIN | VEH | WIN |
| **Cerebral cortex** |  |  |  |  |
| Cingulate | 102 ± 21 | 115 ± 19 | 248 ± 36 | 298 ± 55 |
| Motor |  |  |  |  |
| Layer I-V | 88 ± 16 | 154 ± 17* | 216 ± 41 | 405 ± 72* |
| Layer VI | 169 ± 21 | 154 ± 34 | 353 ± 51 | 354 ± 39 |
| Somatosensory |  |  |  |  |
| Layer I-V | 156 ± 28 | 164 ± 53 | 156 ± 47 | 259 ± 41 |
| Layer VI | 178 ± 21 | 194 ± 38 | 353 ± 66 | 378 ± 96 |
| **Basal ganglia** |  |  |  |  |
| Globus pallidus | 76 ± 33 | 116 ± 75 | 874 ± 115 | 890 ± 103 |
| Striatum | 221 ± 22 | 188 ± 26 | 452 ± 57 | 738 ± 143* |
| **Diencephalon** |  |  |  |  |
| NBM | 129 ± 26 | 195 ± 35 | 231 ± 51 | 277 ± 46 |
| HDB | 139 ± 23 | 244 ± 37* | 230 ± 40 | 257 ± 48 |
| VDB | 247 ± 22 | 250 ± 36 | 224 ± 58 | 613 ± 179* |
| Medial septum | 268 ± 27 | 374 ± 43* | 206 ± 37 | 356 ± 61 |
| **Hippocampus** |  |  |  |  |
| CA1 | 56 ± 17 | 62 ± 17 | 104 ± 27 | 175 ± 22 |
| Oriens | 51 ± 16 | 64 ± 23 | 17 ± 5 | 62 ± 12* |
| Pyramidal | 65 ± 15 | 88 ± 46 | 230 ± 41 | 312 ± 93 |
| Radiatum | 39 ± 7 | 49 ± 8 | 137 ± 23 | 350 ± 87* |
| CA2 | 50 ± 9 | 47 ± 11 | 199 ± 45 | 234 ± 49 |
| CA3 | 48 ± 5 | 36 ± 6 | 276 ± 42 | 425 ± 58* |
| Oriens | 64 ± 23 | 67 ± 18 | 58 ± 19 | 57 ± 12 |
| Pyramidal | 25 ± 19 | 95 ± 29* | 305 ± 110 | 323 ± 130 |
| Radiatum | 34 ± 9 | 30 ± 8 | 74 ± 21 | 184 ± 44* |
| Dentate gyrus | 37 ± 7 | 48 ± 13 | 277 ± 51 | 484 ± 66* |
| Granular | 38 ± 10 | 42 ± 17 | 318 ± 101 | 398 ± 104 |
| Molecular | 40 ± 8 | 34 ± 11 | 174 ± 38 | 308 ± 90 |
| Polimorphic | 27 ± 10 | 27 ± 6 | 324 ± 45 | 338 ± 76 |
| **Amygdala** | 98 ± 18 | 75 ± 31 | 213 ± 59 | 407 ± 87 |

HDB: horizontal diagonal band, NBM: *nucleus basalis magnocellularis*, VDB: vertical diagonal band. Data are expressed as mean ± S.E.M. values from VEH and WIN groups. VEH vs. WIN (\*). Mann-Whitney test, \*p < 0.05.

The [^35^S]GTPγS binding stimulated by WIN55,212-2 was measured in brain areas related to learning and memory to localize and determine the activity of CB_1_ receptors (Fig. 7). The activity mediated by G_i/o_-coupled CB_1_ receptors was increased following the WIN55,212-2 treatment in several areas related to learning and memory processes, such as layers I-V of the motor cortex (VEH *vs*. WIN, p < 0.05; Fig. 7, Table 1), the striatum (VEH *vs*. WIN, p < 0.05; Fig. 7, Table 1), the vertical diagonal band (VEH *vs*. WIN, p < 0.05; Fig. 7, Table 1), the *radiatum* and *oriens* layers of the hippocampus CA1 region (VEH *vs*. WIN, p < 0.05; Fig. 7, Table 1), the hippocampus CA3 region (VEH *vs*. WIN, p < 0.05; Fig. 7, Table 1), the *radiatum* layer of the hippocampus CA3 region (VEH *vs*. WIN, p < 0.05; Fig. 7, Table 1) and the dentate gyrus (VEH *vs*. WIN, p < 0.05; Fig. 7, Table 1).

**Fig. 7.**
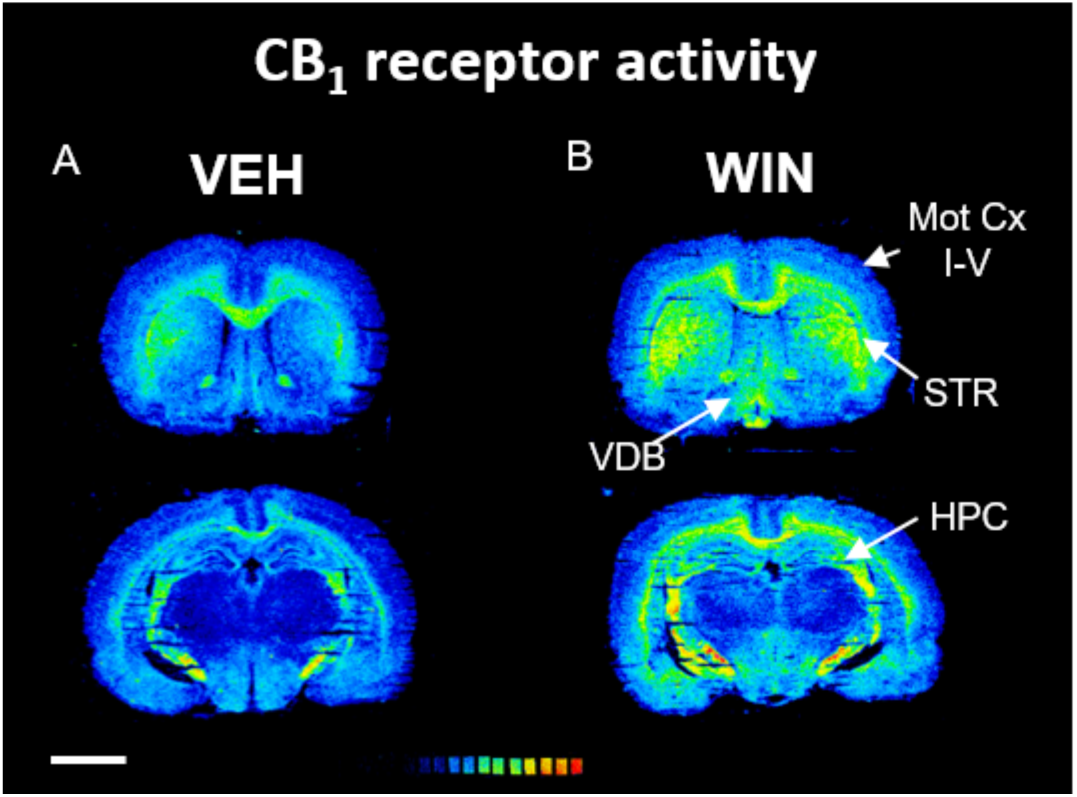
Representative autoradiograms showing the activity of CB1 receptors evoked by WIN55,212-2. Coronal sections corresponding to (A) VEH and (B) WIN groups showing [^35^S]GTPγS binding stimulated by WIN55,212-2 in the vertical diagonal band (VDB), layers I-V of the motor cortex (Mot Cx I-V), the striatum (STR) and the hippocampus (HPC). Scale bar: 4 mm.

The density of CB_1_ receptors was also studied in the same brain areas by [^3^H]CP55,940 binding (Fig. 9). The treatment with the cannabinoid agonist WIN55,212-2 increased CB_1_ receptors density in layer VI of the motor cortex (VEH *vs*. WIN, p < 0.05; Fig. 8, Table 2) and in layer VI of the somatosensory cortex (VEH *vs*. WIN, p < 0.05; Fig. 8, Table 2). In contrast, CB_1_ density decreased in the dentate gyrus (VEH *vs*. WIN, p < 0.05; Fig. 8, Table 2) and in the pyramidal layer of the hippocampus CA1 region (VEH *vs*. WIN, p < 0.05; Fig. 8, Table 2).

**Fig. 8.**
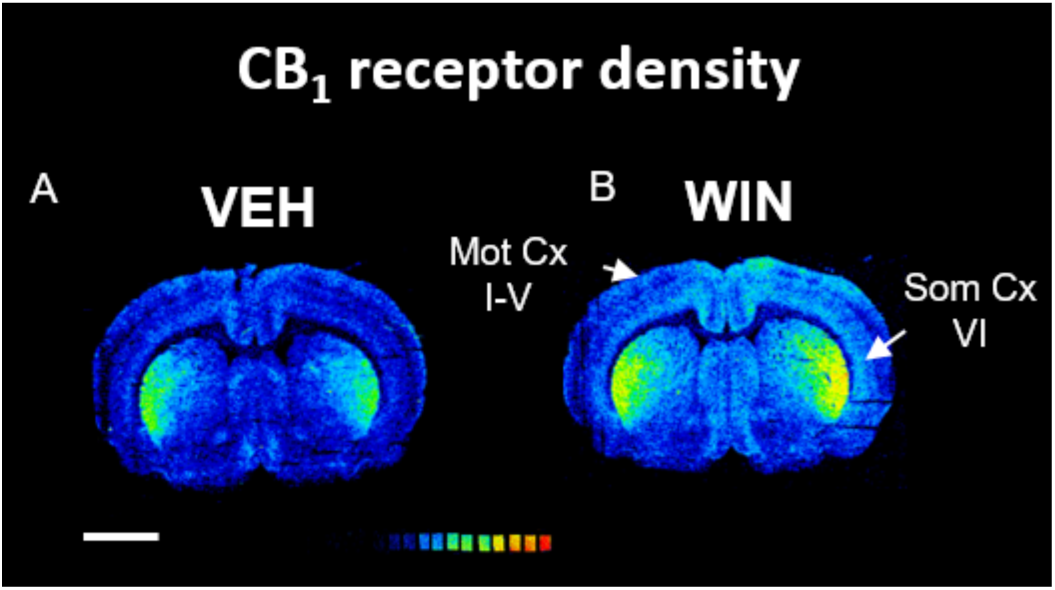
Representative autoradiograms showing the density of CB1 receptors. Coronal sections corresponding to (A) VEH and (B) WIN groups showing [^3^H]CP55,940 binding in layers I-V of the motor cortex (Mot Cx I-V) and in layer VI of the somatosensory cortex. Scale bar: 4 mm.

**Table 2.**
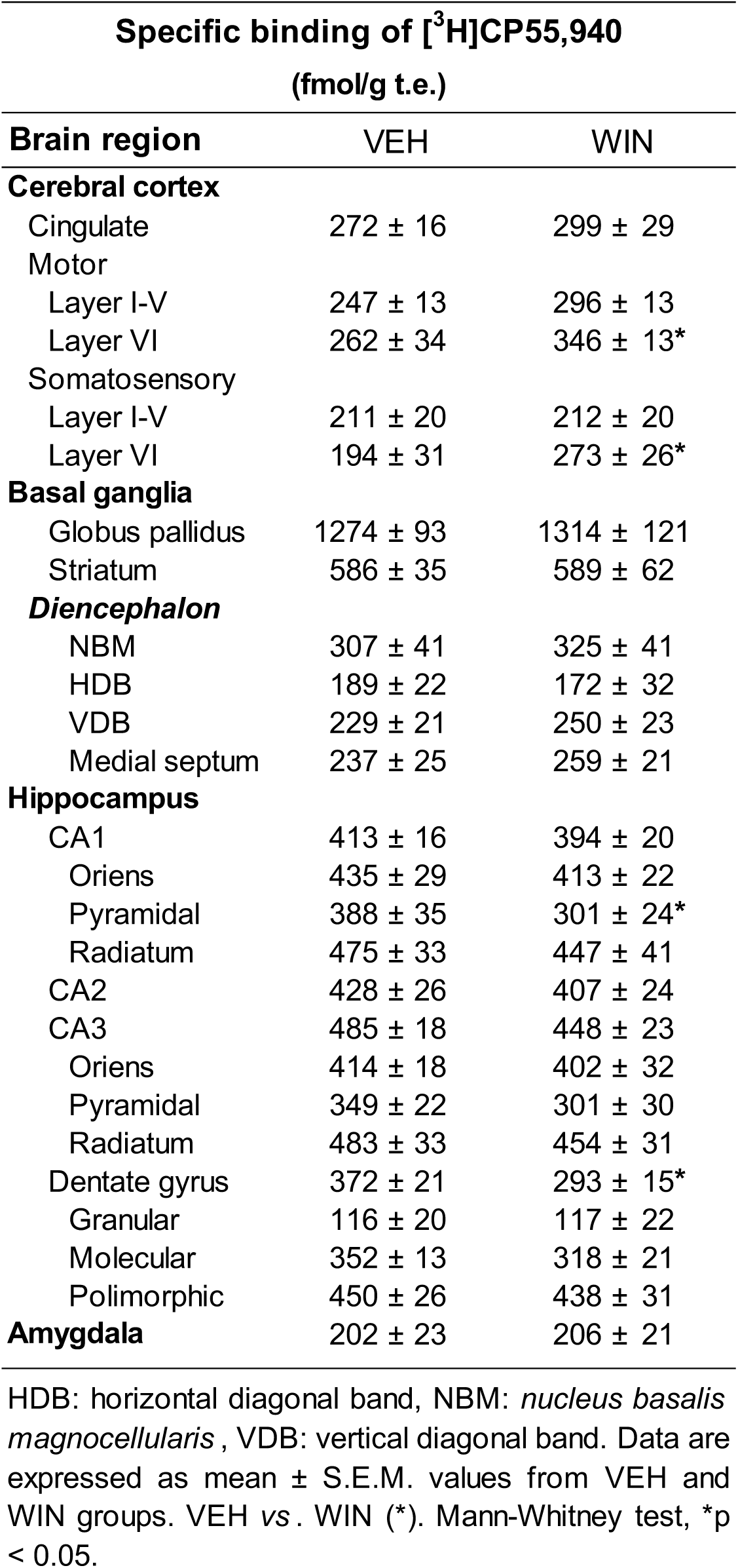
Autorradiographic densities of CB1 receptors expressed in fmol/g t.e., obtained as specific binding of [^3^H]CP55,940, in different brain areas related to learning and memory.

| Specific binding of [<sup>3</sup>H]CP55,940<br>(fmol/g t.e.) |  |  |
| --- | --- | --- |
| Brain region | VEH | WIN |
| **Cerebral cortex** |  |  |
| Cingulate | 272 ± 16 | 299 ± 29 |
| Motor |  |  |
| Layer I-V | 247 ± 13 | 296 ± 13 |
| Layer VI | 262 ± 34 | 346 ± 13* |
| Somatosensory |  |  |
| Layer I-V | 211 ± 20 | 212 ± 20 |
| Layer VI | 194 ± 31 | 273 ± 26* |
| **Basal ganglia** |  |  |
| Globus pallidus | 1274 ± 93 | 1314 ± 121 |
| Striatum | 586 ± 35 | 589 ± 62 |
| **Diencephalon** |  |  |
| NBM | 307 ± 41 | 325 ± 41 |
| HDB | 189 ± 22 | 172 ± 32 |
| VDB | 229 ± 21 | 250 ± 23 |
| Medial septum | 237 ± 25 | 259 ± 21 |
| **Hippocampus** |  |  |
| CA1 | 413 ± 16 | 394 ± 20 |
| Oriens | 435 ± 29 | 413 ± 22 |
| Pyramidal | 388 ± 35 | 301 ± 24* |
| Radiatum | 475 ± 33 | 447 ± 41 |
| CA2 | 428 ± 26 | 407 ± 24 |
| CA3 | 485 ± 18 | 448 ± 23 |
| Oriens | 414 ± 18 | 402 ± 32 |
| Pyramidal | 349 ± 22 | 301 ± 30 |
| Radiatum | 483 ± 33 | 454 ± 31 |
| Dentate gyrus | 372 ± 21 | 293 ± 15* |
| Granular | 116 ± 20 | 117 ± 22 |
| Molecular | 352 ± 13 | 318 ± 21 |
| Polimorphic | 450 ± 26 | 438 ± 31 |
| **Amygdala** | 202 ± 23 | 206 ± 21 |

HDB: horizontal diagonal band, NBM: *nucleus basalis magnocellularis*, VDB: vertical diagonal band. Data are expressed as mean ± S.E.M. values from VEH and WIN groups. VEH vs. WIN (\*). Mann-Whitney test, \*p < 0.05.

## 4. Discussion

Considering the key regulatory role of the eCB system over learning and memory tasks controlled by cholinergic neurotransmission, we have evaluated the effect of a subchronic treatment with potent cannabinoid receptor agonist WIN55,212-2 in preventing transitory amnesia caused by scopolamine. As expected, scopolamine impaired memory in the three learning and memory behavior tests used in the study compared to a saline-treated control group (Malikowska-Racia et al., 2018). The acute pharmacological model of cholinergic blockade with scopolamine is well-characterized and has been applied in the study of numerous anti-amnesic drugs (Akinyemi et al., 2017; El-Khadragy et al., 2014; Marisco et al., 2013). Our results are consistent with data reported in the literature; however, we observed that pretreatment with scopolamine did not significantly disrupt short-term recognition memory in NORT. This observation contrasts with other studies indicating that muscarinic antagonists impair the acquisition and performance of various learned behaviors, including short-term memory (Balderas et al., 2012; Fibiger et al., 1991; Palmer et al., 2016). However, it is consistent with results previously observed in our group, where short-term memory in NORT after a basal cholinergic lesion was not affected, even when baso-cortical cholinergic neurotransmission was greatly diminished (Moreno-Rodríguez et al., 2024). It is also in line with a study reporting that reduced ACh release delays specifically the consolidation of object recognition memory (De Jaeger et al., 2013). Our findings suggest that blocking the cholinergic signaling is crucial for long-term, but not short-term, recognition memory. In our study, WIN55,212-2 alone induced similar cognitive impairment in NORT and PA as scopolamine did, but not in BM. The contradictory effects of a relatively low dose of WIN55,212-2 observed in BM, NORT and PA tests suggest different sensitivities to cannabinoid effects depending on the type of memory assessed, i.e. spatial memory, recognition and contextual memory and aversive or fear-related memory. Previous reports on this matter are somewhat contradictory. For instance, similar doses of WIN55,212-2 facilitated the extinction of contextual fear memory in rats and similar effects were observed in spatial memory (Pamplona et al., 2006). However, different sensitivities to the effects mediated by cannabinoid receptors depending on the type of memory have also been described, with hippocampus-dependent fear memory impaired and hippocampus-independent fear memory remaining unaffected in knockout mice (Li & Kim, 2016). Although the detrimental effect of cannabinoid agonism on memory is well studied and proven, the underlying mechanisms remain unclear.

Here we demonstrated that subchronic administration of low doses of WIN55,212-2 (0.5 mg/kg) prevented the amnesic effects of scopolamine in spatial learning and memory in the BM test, but not in NORT test. Importantly, WIN55,212-2 alone, at this dose, did not induce detrimental effects on memory in BM test, but it did in NORT. In other studies, low doses of THC restored age-related cognitive dysfunction by modulating hippocampus-dependent memory processes. Moreover, the activation of the eCB system in old animals modified proteins and enzymes associated with synaptic connectivity and neuroplasticity specifically in the hippocampus, which contributed to the restoration of certain learning and memory parameters (Bilkei-Gorzo et al., 2017; Sarne et al., 2018). While data suggest cross-state-dependent learning between WIN55,212-2 and scopolamine (Jamali-Raeufy et al., 2011), the differential sensitivity to cannabinoids may be elucidated by exploring the mechanisms through which scopolamine induces memory loss. Scopolamine is a non-selective muscarinic receptor antagonist, which reduces ACh binding to the muscarinic receptors and leads to memory deficits (Blokland, 1995). It mainly affects cholinergic transmission at the basal forebrain, motor cortex, *globus pallidus*, hippocampus, perirhinal cortex and amygdala. However, choline acetyltransferase (ChAT) levels seem to be affected specifically in the hippocampus and amygdala (Hescham et al., 2014; Ray et al., 1992). Scopolamine can decrease cortical ACh levels by 52% and hippocampal ACh levels by 39% (Spignoli et al., 1987). Several studies have shown that cholinesterase inhibitors, which increase ACh levels at the synapse (Pattanashetti et al., 2017; Sadek et al., 2016; Shin et al., 2018) or natural compounds that also increase ACh levels (Lazarova et al., 2024; Teralı et al., 2024), prevent the amnesic effects of scopolamine. However, the effectiveness of memory improvement is dependent on the dose of scopolamine administered (Bejar et al., 1999), suggesting that the levels of ACh in the synaptic cleft are important for the effect of scopolamine. In this sense, it is already described that cannabinoids, including WIN55,212-2, modulate ACh release in the hippocampus and cortex (Gessa et al., 1997; Gessa et al., 1998; Moreno-Rodríguez et al., 2024; Nava et al., 2001;Tzavara et al., 2003).

WIN55,212-2 can also mitigate the toxicity induced on the cholinergic system following acetylcholinesterase inhibition (Nallapaneni et al., 2006; Nallapaneni et al., 2008). Therefore, we hypothesise that the WIN55,212-2-mediated increase in ACh could potentially compensate for the acute reduction in cortical and hippocampal ACh levels induced by scopolamine (Spignoli et al., 1987), as we have previously described in cortical areas (Moreno-Rodríguez et al., 2024). In addition, WIN55,212-2 may induce memory deficits itself due to an excessive increase in ACh levels, as recent evidence suggests (Huang et al., 2022), among other mechanisms. However, the mechanism by which the same doses of cannabinoid agonists protect against scopolamine-induced cognitive impairment in the BM while showing no effect on NORT remains unclear. While BM and NORT are both hippocampus-dependent tests, BM may require higher modulation of ACh to facilitate precise navigation and spatial memory, whereas NORT may be more sensitive to moderate levels of ACh for the effective processing of visual information and recognition memory (Antunes & Biala, 2012; Haam & Yakel, 2017; Pitts, 2018). The levels of ACh required for recovery from scopolamine-induced impairment in the BM may, thus, be excessive for NORT.

Regarding the third test performed in this study, PA, the deleterious effects of cannabinoids on aversive memory are well known and have been previously described (Jamali-Raeufy et al., 2011). However, cannabinoid compounds also affect the ascending and descending pathways that regulate pain perception (Starowicz & Finn, 2017). This is relevant, considering the aversive stimulus (a mild foot shock) inherent to this task (Bengoetxea de Tena et al., 2022). Indeed, our results indicate that WIN55,212-2 at this dose altered pain sensitivity in rats, likely influencing the learning and memory outcomes in the PA test.

Finally, our subchronic treatment with WIN55,212-2 also modified motor activity. The animals treated with WIN55,212-2 moved more slowly compared to those treated with vehicle. However, the unchanged total path length during training in BM test suggests that this reduction in movement did not affect spatial acquisition. Cholinergic denervation of the forebrain enhances locomotor activity responses associated with dopaminergic activity (Mattsson et al., 2002). Additionally, ACh release in the striatum, hippocampus, and frontal cortex correlates with locomotion (Day et al., 1991), suggesting that the status of the cholinergic system affects motor activity. The opposite effect, a reduction in locomotor activity following WIN55,212-2 administration, has also been described (Compton et al., 1992). The capacity of cannabinoids to modulate motor activity, as well as cognition, may be attributed, at least partly, to alterations in cortical cholinergic neurotransmission. The results from the autoradiographic assays in rats that completed the BM test support this hypothesis. Following the WIN55,212-2 treatment, the activity of muscarinic M_2_/M_4_ receptors specifically increased in parts of the motor cortex and the hippocampus. These results apparently contrast with others that show WIN55,212-2 treatment leading to a decrease in M_2_ receptor densities (Schulte et al., 2012) and a down-regulation of M_2_/M_4_ receptors in animals with high levels of ACh in the hippocampus (du Bois et al., 2005). In specific layers of these same areas, the density and activity of CB_1_ receptors was also modulated following WIN55,212-2 administration. As previously mentioned, the activation of these cannabinoid receptors trigger an increase in ACh levels, by mobilizing specific choline-containing phospholipids which could explain the enhanced muscarinic receptor activity (Moreno-Rodríguez et al., 2024). We hypothesise that this increase was enough to counteract the effects of scopolamine in BM test, but it was excessive for NORT. Our group has previously described the positive effects of WIN55,212-2 in the BM. We demonstrated that WIN55,212-2 increased ACh levels in a dose-dependent manner in the cortex and restored cortical cholinergic neurotransmission after a basal forebrain cholinergic lesion (Moreno-Rodríguez et al., 2024). Based on the results from this study, where the same protocol for WIN55,212-2 administration and performance in BM was followed, we propose a similar mechanism. Since scopolamine induces an acute reduction of ACh levels, the previously elevated ACh levels caused by WIN55,212-2 treatment prevent scopolamine from exerting deleterious cognitive effects.

## 5. Conclusions

Overall, these results suggest that, at low doses, a treatment with WIN55,212-2 can prevent the amnesic effects induced by scopolamine in a spatial learning and memory test like BM, but not in a recognition memory test, such as NORT. The specific mechanisms underlying this effect remain elusive, but results suggest a differential modulation of the crosstalk between the eCB and the cholinergic systems depending on the type of memory assessed in each test. More precisely, the subchronic activation of CB_1_ receptors potentially increased the cholinergic tone in key cortical and hippocampal areas, just enough to overcome the scopolamine-induced cholinergic deficit in BM test. The prospective clinical application of this promising experimental data could be related to the modulation of the eCB system for the treatment of dementias associated with a cholinergic deficit, like is the case of AD.

## Funding

This research was financially supported by grants from the Basque Government to the “Neurochemistry and Neurodegeneration” consolidated research group (IT975-16 and IT1454-22 to R.R-P), by Instituto de Salud Carlos III, co-funded by European Regional Development Fund “A way to make Europe” (PI20/00153 to R.R-P) and by BIOEF funded by Eitb Maratoia (BIO22/ALZ/010 to R.R-P). I.B.d.T was the recipient of an Investigo fellowship funded by the European Union Next Generation. G.P-C was the recipient of a University of the Basque Country predoctoral fellowship.

## CRediT authorship contribution statement

Marta Moreno-Rodríguez and Iker Bengoetxea de Tena contributed equally to this work and share first-authorship. **Marta Moreno-Rodríguez:** Conceptualization, Validation, Formal analysis, Investigation, Writing - Original Draft, Writing - Review & Editing, Visualization. **Iker Bengoetxea de Tena:** Conceptualization, Validation, Formal analysis, Investigation, Writing - Original Draft, Writing - Review & Editing, Visualization. **Jonatan Martínez-Gardeazabal:** Investigation, Writing - Review & Editing. **Gorka Pereira-Castelo:** Investigation, Writing - Review & Editing. **Alberto Llorente-Ovejero:** Investigation, Writing - Review & Editing. **Iván Manuel:** Investigation, Writing - Review & Editing. **Rafael Rodríguez-Puertas:** Conceptualization, Resources, Writing - Review & Editing, Supervision, Project administration, Funding acquisition.

## Declaration of competing interest

The authors declare that the following Spanish patent related to the present work has been registered: *Tratamiento de la demencia con agonistas cannabinoides. Spain. 02-03-2017. University of the Basque Country. ES2638057*.

## Data availability

The data that support the findings of this study are available from the corresponding author upon reasonable request.

